# Crossmodal pitch-luminance association in tortoises

**DOI:** 10.1101/2025.01.24.634651

**Authors:** Maria Loconsole, Beatrice Malaman, Gionata Stancher, Elisabetta Versace

## Abstract

If you think that high-pitched sounds match bright colours more than dark ones, you are not alone. Spontaneous associations between different sensory modalities —crossmodal associations— are widely observed in humans. More recently, crossmodal associations have been reported in chimpanzees, monkeys, dogs, chickens, and tortoises, indicating a general cognitive strategy that reflects natural correlations in the environment or similarities in the architecture of the nervous system. Alternatively, or complementarily, crossmodal associations may arise from learning of species-specific occurrences. Recent research on primates has tentatively linked human pitch-luminance associations to the evolution of language. While humans and chimpanzees spontaneously match a high-pitched sound with a white shape and a low-pitched sound with a black shape, baboons and chickens do not exhibit any preferential association. Identifying spontaneous pitch-luminance associations in species that do not rely on vocal communication, such as the solitary Hermann’s tortoises, would support the idea of a language-independent phenomenon. To test this idea, we studied tortoises for pitch-luminance associations in a spontaneous food-searching task. After hearing a high-pitched (700 Hz) vs low-pitched (450 Hz) sound, animals could choose to search for food behind either a light- or dark-coloured wall. When hearing the high-pitched sound, tortoises chose the white stimulus, whereas when hearing the low-pitched sound, they chose the black stimulus, similarly to what humans and chimpanzees do. These results demonstrate crossmodal pitch-luminance associations in tortoises. While showing this commonality in the perceptual organization of phylogenetically distant species, we shift the question on whether pitch-luminance associations reflect homology or convergent evolution.

## Introduction

Despite sensory receptors being segregated, the perception across sensory modalities is so closely connected that hearing a high-pitched sound spontaneously prompts the idea of a small and light-coloured visual stimulus, whereas hearing a low-pitched sound activates the idea of a large and dark-coloured object ^[1]^. These crossmodal correspondences between different sensory modalities have also been reported in non-human species, e.g., chimpanzees ^[2]^, rhesus monkeys ^[3]^, dogs ^[4,5]^, chickens ^[6]^, and tortoises ^[7]^. Altogether, these findings suggest that crossmodal associations may represent a widespread cognitive strategy, potentially rooted in the ability to detect and interpret natural correlations across sensory modalities or reflecting similarities in the neural organization/coding ^[1,8]^. This shared capacity could reflect an evolutionary advantage in processing multisensory information, enhancing perception and decision-making across diverse ecological contexts. Alternatively, or complementarily, crossmodal associations may stem from learning processes and species-specific cognitive skills shaped by ecological and social contexts ^[9]^. Recent findings in primates have proposed a potential link between human pitch-luminance associations and the evolution of language, suggesting that such crossmodal pairings might be influenced by the unique communicative and cognitive demands of humans and closely related species. Humans ^[2,10]^ and chimpanzees ^[2]^ reliably associate high-pitched sounds with white shapes and low-pitched sounds with black shapes. When tested in a categorisation task of white and black shapes, subjects were faster in responding when hearing a background sound congruent to the pitch-luminance association (i.e., high-pitched when responding to white, and low-pitched when responding to black). In contrast, baboons tested in the same task did not show any consistent preference. Moreover, three-day old domestic chickens ^[11]^ tested in a similar task, where they had to choose between a black or a white panel to locate a food reward, did not show any facilitation related to high- or low-pitched background sounds, indicating that this association may not be universal across all species. Novel evidence of pitch-luminance associations in non-primate species that do not depend on vocal communication could provide crucial insights into the origins of this phenomenon. Specifically, studying solitary species like Hermann’s tortoises offers an opportunity to test whether such associations are intrinsic and independent of language or social learning. To explore this, we tested 18 tortoises (*Testudo hermanni*) in a food-searching task comprising 16 consecutive trials where the animals could spontaneously choose to search food behind either a white (high luminance) or a black (low luminance) wall (**Fig. 1**). During each trial, the same background sound was played simultaneously behind the two doors, either a high-pitched (700 Hz) or low-pitched (450 Hz) sound. In these unrewarded trials, tortoises spontaneously chose the white door when hearing the high pitch sound, and the black door when hearing a low pitch sound (450 Hz). This result is similar to what previously observed in humans and chimpanzees. Together with mounting evidence of crossmodal associations in different taxa ^[4–6]^, data from a reptile suggest the idea of the widespread presence of spontaneous crossmodal associations as a language-independent and predisposed mechanism.

**Fig. 1.**
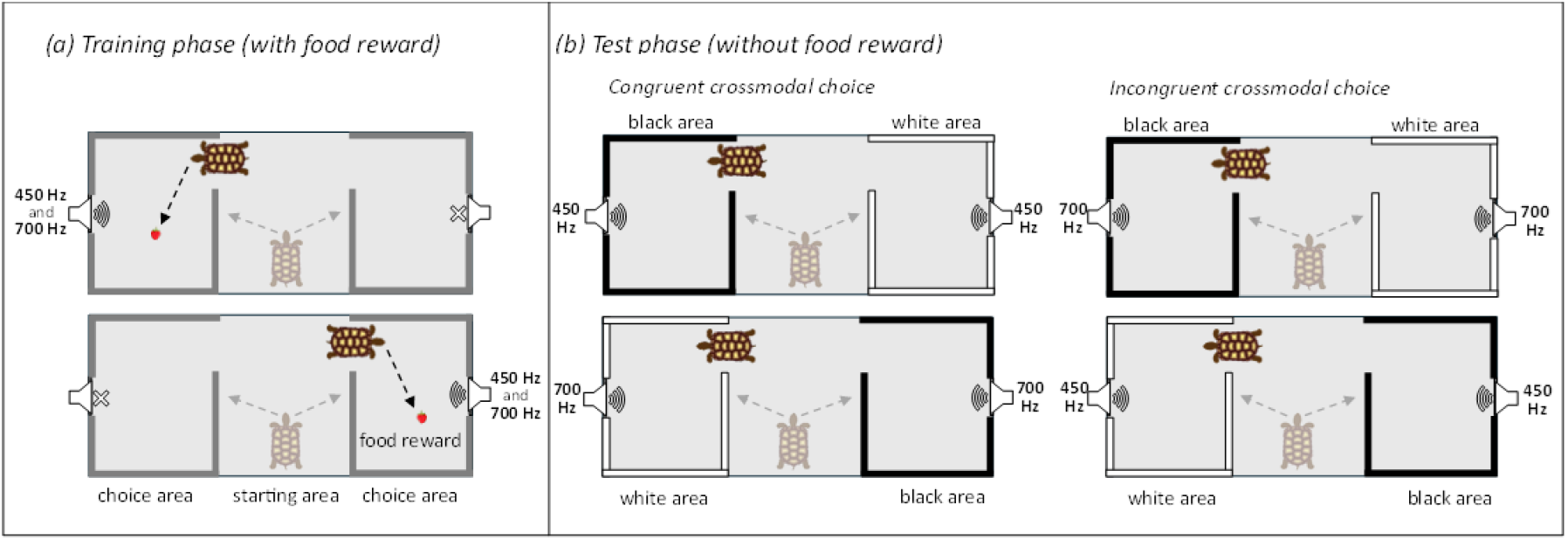
The experimental arena and procedures. The arena is divided into three separate areas: the subject is placed into the central starting area and is free to enter each of the two side areas (choice areas) via a small opening on the side. (a) In each training trial, a piece of strawberry was placed close to the playing loudspeaker. The loudspeaker alternated high (700 Hz) and low (450 Hz) pitch sounds, each lasting 10second. The arena was dyed all grey. At training subjects learnt to approach the source of the sound to find the reward. (b) At test, tortoises received no reward. Both speakers played the same pitch sound (either high or low). Each choice area can be identified by the colour of the walls (either white or black). We hypothesized a preference for the white area in the high-pitch sound condition and a preference for the black area in the low-pitch sound condition (congruent crossmodal choice) rather than the opposite (incongruent crossmodal choice).

## Materials and methods

### Subjects

We tested 18 adult male tortoises (*Testudo hermanni*) aging between 15 and 30 years. Tortoises were housed at Sperimentarea (Rovereto Civic Museum Foundation, IT) in a semi-free environment, consisting of two open fields of approximately 10 × 12 meters each, with natural vegetation delimited by metal fences, where the animals lived in same-sex groups with free access to food and water. These animals are usually brought to Sperimentarea by the local authorities, either after being confiscated or found abandoned. Since tortoises are not endemic in this area, we expect all or most of them to be born in captivity.

The experimental subjects were temporarily housed in a separate area within the main facility, where they had free access to fresh water and were regularly fed lettuce and herbs. To avoid disturbing egg-laying in females, only male tortoises were included in the study. To avoid environmental confounds, animals were tested individually in a shed separated from their living environment. The experiments were carried out in July-August 2024, during the period of highest activity (tortoises fall into hibernation approximately from November to April).

This study complied with all applicable national and European laws concerning the use of animals in research. All the employed procedures were examined and approved by the Rovereto Civic Museum Foundation Ethical Review Committee (Prot. n. 0000132 dd. 12.04.2023).

### Experimental arena and stimuli

The experimental arena consisted of a wooden corridor (50 cm high) divided into three separate areas: a central area (50 × 50 cm) that served as the animal’s starting point, and two choice areas (35 × 50 cm), each accessible through a small opening in the walls (25 × 10) on either side of the starting area. On each of the two short sides of the arena we located a loudspeaker (Ex!treme Flat Panel Speaker with Amplifier system, model P-188) to play the auditory stimuli. Mounted above the arena there was a 400 W halogen lamp and a video camera (Microsoft HD) for recording the tests. Auditory stimuli consisted of a 10-second high-pitch (700 Hz) sound, and a 10-second low-pitch (450 Hz) sound, prepared by using the software Audacity, setting an amplitude of 65 dB. A previous work on pitch-size correspondence ^[7]^ that employed a similar methodology showed that tortoises are sensitive and responsive to these two frequencies.

### Training

The experiment started with a familiarisation and training procedure aimed at acquainting the subjects to a neutral grey arena and teach them to reach the source of the sound to find a palatable reward, a small piece of strawberry. In this stage, the arena’s floor and walls were lined with 290 g opaque grey contact paper (opaque polyvynil cloryde). Each tortoise underwent several training sessions, from a minimum of 3 to a maximum of 6. Each session lasted approximately 30 minutes and consisted of a series of consecutive trials. If the subject reached a maximum of 24 trials, or exceeded 30 minutes of training, the session ended and the subject was allowed to rest for minimum 2 hours. In each trial, only one speaker was active, playing an alternation of the high- and low-pitch sounds spaced out by 1 s silence. The position of the playing speaker was counterbalanced between trials.

Training animals to the presence of food hidden behind a door, close to the sound source was achieved through a shaping procedure. The animal was placed into a starting box in the centre of the empty arena with the strawberry close to the playing loudspeaker. Next, the two central walls (each with a small lateral opening that allowed access behind the wall) were added, creating three distinct areas. At this stage, the strawberry was still visible at the entrance of the correct opening. Finally, the strawberry was hidden behind the wall, so the tortoises could only rely on the auditory cue to locate the correct area to enter. The position of the reward (left or right opening with respect to the tortoise position) was counterbalance between trails. The first/second sound heard (alternation of low-high, or high-low pitch sounds) was counterbalanced between trials.

When the subjects entered the correct area (area with sound and strawberry), they were allowed to eat the food reward. Then they were removed from the arena and a new trial was started. When the subjects entered the incorrect area, they were immediately removed from the arena without the possibility to enter the other area, and a new trial was started.

Each subject moved to the next step of the shaping procedure when they responded correctly to six consecutive trials out of eight, within three minutes per trial. If this criterion was not met, the last step of the shaping procedure was repeated.

### Test

The test was always conducted in a single, separate session from the training. Before undergoing the test, each subject went through a refresh phase consisting of four additional training trials. Then, each animal underwent 16 consecutive test trials.

During test, the starting area remained the same as in training, but the adjacent walls were of different colours. One was white (170 hue, 0 saturation, 255 lightness), with the corresponding opening leading to a white area, and the other wall was black (170 hue, 0 saturation, 0 lightness), with the corresponding opening leading to a black area. During testing, both speakers were active, meaning that the source of the sound was no longer a reliable cue for locating the food reward (**Fig. 1**). For each trial, the speakers played a repetition of the same sound, either the high-pitch, or the low-pitch, with 1 second of silence between each repetition. The position of the white and black walls (i.e., left or right), as well as the pitch of the sound, were pseudo-randomly alternated between trials. No food reward was present during test to rule out possible olfactory confounding.

We hypothesised that if tortoises rely on predisposed crossmodal associations to solve the test, they should choose the white area more often when hearing the high-pitch sound, and the black one when hearing the low pitch-sound ^[2]^.

### Quantification and statistical analysis

The raw data generated during the study are available as supplementary material (**S1**). Data were analysed using R 4.4.0 ^[12]^. Alpha was set to 0.05. We used a generalised linear mixed-effect model (R package: lme4 ^[13]^) with the dependent variable being binomial: 1 = congruent association (that is, between high pitch and white, and low pitch and black, as reported in humans (2)), 0 = incongruent association between pitch and luminance. We ran an Akaike information criterion-based model selection to determine the minimum adequate model, including as predictors the trial (from 1 to 16), the pitch (high or low), the position of the white area (left or right), and their interactions. We run an analysis on the selected model using the R package emmeans ^[14]^. The graph was generated using ggplot2 ^[15]^.

## Results

Analysis of the minimum adequate model (based on the Akaike Information Criterion – AIC) revealed that tortoises relied on a congruent association for which they chose more often the white door when hearing the high pitch sound, and the black door when hearing the low pitch one (prob(congruent association) = 0.612, SE = 0.03, z = 3.755, p < 0.001). This effect remains consistent across trials. Regarding a potential effect of experience with unrewarded trials during the test, and the issue of plasticity of crossmodal associations, we observed a trend for a decrease in pitch-luminance association during the test (**Fig. 2**), although not statistically significant (effect of trial: β = −0.436; SE = 0.026; z = −1.647; p = 0.1).

**Fig. 2.**
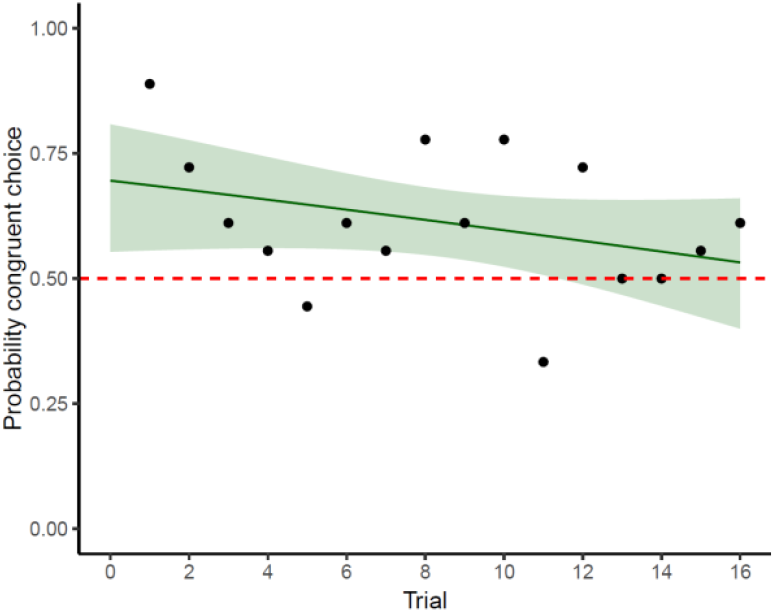
Probability of displaying the hypothesized crossmodal association among 16 testing trials when considering the door chosen by the subject. Each point represents group performance; the shaded area represents the confidence interval; the dashed red line represents chance level (i.e., probability of 0.5).

## Discussion

Our results show that Hermann’s tortoises spontaneously associate a high-pitched sound with brightness (white) and a low-pitched sound with darkness (black), similar to what humans ^[2,10]^ and chimpanzees ^[2]^ do. The presence of this pitch-luminance correspondence in a reptile suggests that the ability to link auditory and visual stimuli may have deep evolutionary roots, potentially serving adaptive functions across diverse ecological contexts or emerging from general advantages of sensory integration ^[16]^. For instance, crossmodal associations may stem from shared processing of intensity features, with brightness and loudness potentially encoded by the same brain structures dedicated to magnitude ^[16,17]^. The fact that some human languages use terms like ‘high’ and ‘low’ to describe both pitch and luminance could reflect an inherent predisposition to link these modalities, rather than language itself being the origin of the association. Together with mounting evidence of spontaneous crossmodal correspondences in different taxa ^[2–7]^, our results support the idea that crossmodal associations are in place as a fundamental building block of knowledge from early life ^[18,19]^, and challenge the idea that such associations could derive from language-related ecological pressures. However, it remains unclear why other species, such as baboons and chickens, did not show similar crossmodal associations in comparable tests. Methodological factors and previous experience may explain these discrepancies. For example, chicks might have been overly exposed to high pitches and brightness during rearing, which could have influenced their responses. Additionally, the auditory stimuli used for testing chicks were the same as those for primates and may not have been ecologically relevant to birds, whose most sensitive auditory range is between 600–2500 Hz ^[11]^. In the case of baboons, it has been suggested that attention issues or difficulty discriminating the auditory stimuli, coupled with their extensive experience with similar training protocols and visual tasks, could have affected subjects’ performance ^[10]^. This is consistent with the notion of spontaneous associations updating with experience ^[7,11]^. Previous studies in which animals were tested for other instances of crossmodal associations in extinction ^[6,7]^ found that the effect diminished over repeated unrewarded trials, eventually reaching chance level. This is evidence that predisposed multimodal associations are flexible and subject to plasticity. Such flexibility is crucial for this mechanism to remain advantageous, enabling the subject to switch to a different strategy if their prior proves unhelpful ^[7]^. or based on frequency of exposure ^[11,20]^. It is also possible that pitch-luminance crossmodal associations genuinely do not exist in these species ^[10]^. Directly retesting these species with paradigms adapted to their sensory and ecological needs will be cricial to disentangle the effect of experience and methodological influences from true interspecies variability ^[21]^. Understanding the phylogenetic development of this phenomenon is essential for a deeper comprehension of sensory integration, cognitive evolution, and the universality of crossmodal perception across different species.

## Supporting information

raw data

## Acknowledgments

We wish to thank all the staff from Sperimentarea for the help in building the experimental arenas and animal care, and the Rovereto Civic Museum Foundation for providing the facilities to carry out this research. Maria Loconsole’s work is funded by the European Union – NextGenerationEU and by the University of Padua under the 2023 STARS Grants@Unipd programme (project CROSS - Comparative Research Of Sound Symbolism). Beatrice Malaman is founded by the Italian Ministry of Education and Research through the Research Project of National Relevance (PRIN) – 2022 Prot. 2022TYX52H. Elisabetta Versace was supported by Leverhulme Trust research grant RPG-2020-287.

## Author Contributions

Conceptualisation, M.L., E.V.; Methodology, M.L., B.M., G.S:, E.V.; Validation M.L., B.M., E.V.; Formal Analysis, M.L., E.V.; Investigation, M.L., B.M.; Resources, M.L., G.S., E.V.; Data Curation, M.L., B.M.; Writing – Original Draft, M.L.; Writing – Review & Editing, B.M., G.S:, E.V.; Visualization, M.L., B.M., E.V.; Supervision, M.L., G.S:, E.V.; Project Administration, M.L., G.S:, E.V.; Funding Acquisition, M.L., E.V.

## Competing Interest Statement

The authors declare no competing interest

## Notes

### Competing Interest Statement

The authors have declared no competing interest.

